# The interactive effect of sustained sleep restriction and resistance exercise on skeletal muscle transcriptomics in young females

**DOI:** 10.1101/2024.02.15.580577

**Authors:** Olivia E. Knowles, Megan Soria, Nicholas J. Saner, Adam J. Trewin, Sarah E. Alexander, Spencer S.H. Roberts, Danielle Hiam, Andrew Garnham, Eric J. Drinkwater, Brad Aisbett, Séverine Lamon

## Abstract

**Introduction:** Both sleep loss and exercise regulate gene expression in skeletal muscle, yet little is known about how the interaction of these stressors affects the transcriptome. The aim of this study was to investigate the effect of nine nights of sleep restriction, with repeated resistance exercise (REx) sessions, on the skeletal muscle transcriptome of young, trained females.

**Methods:** Ten healthy females aged 18-35 years undertook a randomised cross-over study of nine nights’ sleep restriction (SR; 5-h time in bed) and normal sleep (NS; ≥7 h time in bed) with a minimum 6-week washout. Participants completed four REx sessions per condition (day 3, 5, 7 and 9). Muscle biopsies were collected both pre- and post-REx on days 3 and 9. Gene and protein expression were assessed by RNA sequencing and Western Blot, respectively.

**Results:** Three or nine nights of sleep restriction had no effect on the muscle transcriptome independently of exercise. However, close to 3000 transcripts were differentially regulated (FDR < 0.05) 48 h post the completion of three resistance exercise sessions in both NS and SR conditions. Only 39% of downregulated and 18% of upregulated genes were common between both conditions, indicating a moderating effect of sleep restriction on the response to exercise.

**Conclusion:** Sleep restriction and resistance exercise interacted to alter the enrichment of skeletal muscle transcriptomic pathways in young, resistance-trained females. Performing exercise when sleep restricted may not provide the same adaptive response for individuals as if they were fully rested.

## Introduction

Adults sleep for approximately one third of their lives, yet up to 40% of the population is failing to obtain adequate sleep (1). Inadequate sleep incorporates both insufficient duration and quality of sleep (2). The negative consequences of chronic exposure to sleep restriction (i.e. <7 hours per night) are becoming increasingly recognised (3), including a loss of muscle mass and function that may increase the likelihood of sarcopenia (4) and other metabolic diseases such as type 2 diabetes and obesity (3, 5). The maintenance of skeletal muscle mass and strength in the face of inadequate sleep is therefore critical to ensuring optimal immune health, physical capacity and metabolism.

Skeletal muscle mass is maintained by a balance of protein synthesis and protein degradation. Sleep deprivation (i.e. 24 h wakefulness) (6) and sleep restriction (i.e. shortened sleep duration) (7) blunt skeletal muscle protein synthesis in young males, yet the mechanisms by which this occurs remain equivocal. In rodent models, some molecular markers of muscle protein synthesis (MPS; e.g. Akt Ser 473 phosphorylation status) are suppressed and molecular markers of muscle protein degradation (MPD; e.g. FOXO3 phosphorylation) are elevated with 96 h of paradoxical sleep deprivation (8, 9). In human males, these markers have however shown no changes with sleep restriction or deprivation (6, 7). Skeletal muscle function and metabolism are further regulated by an internal circadian clock. Some authors (10) but not others (6) have reported changes in the expression of circadian clock genes with inadequate sleep. At the muscle transcriptome level, 24 h sleep deprivation (10) but not five repeated nights of sleep restriction (4 h sleep per night) (11), alter the expression of individual genes associated with the regulation of circadian clock and protein metabolism pathways in males.

The duration of sleep deprivation or restriction, physical activity status, timing of muscle sample collection, and sex (12) are factors that may impact transcriptional changes in the muscle. In addition, while being more likely to experience sleep problems than males (13), females are also underrepresented in the field of sleep (14) and muscle physiology (15) research. Despite proposed sex-specific mechanisms in how sleep regulates muscle protein and gene expression (6, 16, 17), our current knowledge remains predominantly based on male data.

Exercise promotes wide-ranging health benefits, and for population groups such as athletes, military service members and shiftworkers, is often performed concurrently to the stressor of sleep loss. Saner et al. (7) showed that high-intensity interval exercise may counteract the adverse effects of sleep restriction in males by maintaining myofibrillar protein synthesis rates at baseline levels. Resistance exercise is an even more potent stimulus to promote MPS (18) and may constitute a better intervention to maintain muscle mass and function in sleep-restricted populations (19). Resistance exercise is known to induce a large, but transient transcriptional response, which is thought to underpin the associated increases in strength and muscle mass (20). We have previously shown that sleep restriction impairs resistance exercise performance (21), but how sleep restriction may impact the muscle molecular response to resistance exercise over time is unknown.

The aim of this study was to investigate the effect of nine nights of sleep restriction, with repeated resistance exercise, on the muscle transcriptome of young, trained females. A better understanding of the interaction between inadequate sleep and muscle adaptation in females may assist in implementing strategies to attenuate skeletal muscle health-related diseases associated with inadequate sleep.

## METHODS

### Participants

Fourteen healthy, resistance trained (defined as having been performing structured resistance exercise at least twice per week for the previous six months) females aged 18-35 years were recruited to participate in this study. Four participants withdrew for personal reasons; therefore, ten participants were included in the final sample. At the time of study design, no similar human studies examining sleep restriction and molecular markers of muscle adaptation (to resistance exercise) existed on which to base sample size calculations. Therefore, the power calculation was determined for the primary outcome measure of the broader study only (resistance exercise volume load), and all other measures were included as secondary analysis (21).

This study is a sub-component of a broader study investigating the effect of sustained sleep restriction on muscle strength performance, and the participant inclusion and exclusion criteria have been described elsewhere (21). All participants regularly slept >7 h per night and had a ‘moderately morning’ or ‘neither’ chronotype classification, as determined using a combination of actigraphy and self-report measures, and the morningness-eveningness questionnaire (22), respectively. All participants met an average daily energy contribution of protein within 15-25% of their total macronutrient intake, as assessed by ASA24® dietary recalls (23), and did not take a protein supplement. All participants were eumenorrheic (i.e., menstruation occurs consistently on a 21- to 35-day cycle). While recent research suggests a minimal effect of menstrual cycle phase on muscle strength (24), we still aimed to minimize any potential confounding impact (25) on any other muscle outcome, and a detailed account of the strategy used to minimize the effects of menstrual cycle fluctuations, participant ovarian hormone data and menstrual phase identification can be found elsewhere (21).

### Study protocol overview

This study was approved by the Deakin University Human Research Ethics Committee (DUHREC 2017-305) and conducted in accordance with *The Declaration of Helsinki* (1964) and its later amendments. The pre-condition, sleep restriction (SR) condition and normal sleep (NS) condition protocols are described in detail elsewhere (21). Briefly, participants attended the laboratory on two occasions prior to the experimental conditions to 1) provide written informed consent, undertake a baseline blood sample and 3-repetition maximum strength measurements and, 2) complete a resistance exercise session familiarisation. A cross-over design was implemented, and participants were randomly allocated to the SR condition (followed by a minimum 12-week washout period) or NS condition (followed by a minimum 6-week washout period) first. Caffeine and alcohol intake were ceased 48-h prior to and during each condition.

During the SR condition, participants spent nine consecutive nights at the sleep laboratory with a 5-h sleep opportunity from 0100 h to 0600 h each night (Figure 1). Lighting was dimmed <100 lux (Digitech Lux Meter, Reduction Revolution, Sydney, NSW) between 2100 h and 0700 h and electronic devices removed between 2300 h to 0630 h. When electronic devices were removed, participants played card or board games to ensure they stayed awake prior to lights out. At 1500 h on days 3, 5, 7 and 9, participants completed a 45-min supervised resistance exercise session designed to replicate real-world training and stimulate muscle protein turnover. On days 3 and 9, muscle biopsies were performed immediately prior to and 1-h post resistance exercise to capture the maximum activation of MPS (26). Muscle samples were obtained under local anaesthesia (2 mL per biopsy site of 1% Lidoocaine, without epinephrine) from the belly of the vastus lateralis muscle using the percutaneous biopsy technique (27). Upon collection, muscle samples were immediately frozen in liquid nitrogen, and stored at –80°C until subsequent analysis. Blood samples were collected pre- and post-exercise to confirm menstrual cycle phase and hormone responses, described elsewhere (21). On days 5 and 7, resistance exercise sessions were performed with blood samples collected only. Data from participant’s ASA24® dietary recalls (23) was used to inform daily macronutrient consumption and energy intake. Participants were instructed to maintain their usual diet during the NS condition and were then provided with individualised macronutrient energy-matched meals that corresponded to their usual intake during the SR condition. This strategy was implemented to avoid a change in diet influencing muscle protein turnover (28). Baseline muscle biopsies were performed at least 2 h after participant’s lunchtime meal and remained fasted until the second biopsy. Outside of the resistance exercise sessions and sleep periods, participants were permitted to study, read, talk with staff, and walk at a low intensity.

**Figure 1.**
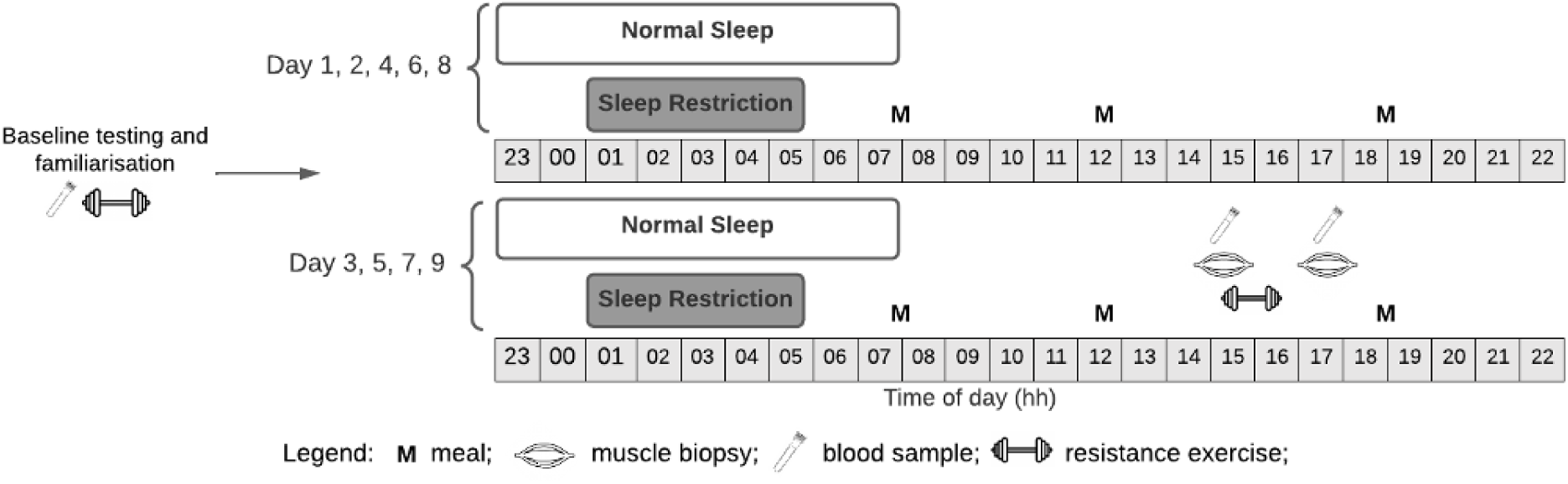
Overview of the study protocol. Sleep was monitored for six days prior to each condition. The sleep opportunity period was between 2200 h and 0800 h for Normal Sleep (NS) and between 0100 h and 0600 h for Sleep Restriction (SR). A 6-12 week washout was implemented between conditions.

During the NS condition, participants remained in their homes and attended the laboratory only on days 3, 5, 7 and 9 to complete the blood sampling, muscle biopsies and resistance exercise. Both conditions were conducted in the same season to equalise the duration of natural light exposure during the day. Participants were instructed to sleep normally with sleep time permitted between 2200 h and 0800 h to achieve similar living and sleeping conditions in the NS condition as compared to the SR condition (i.e. lighting dimmed for 30min prior to sleep and upon waking, similar temperature, low noise during sleep periods and low levels of low intensity physical activity permitted during waking periods). Sleep diaries and ASA24® dietary recalls were completed from home, with reminders sent via text message following mealtimes.

## Measures

### Sleep

Activity monitors (Actical Z MiniMitter, Philips Respironics, Bend, OR) were used in conjunction with sleep diaries to assess sleep. Monitors were worn for six days prior to, and throughout each experimental condition. These were worn 24 h per day on the non-dominant wrist, except when showering or bathing. Activity counts were recorded in one-minute epochs via an embedded piezoelectric accelerometer that records movements in all planes (29). Raw data were downloaded following completion of each condition using a device specific interface unit (ActiReader, Philips Respironics, Bend, OR). A propriety algorithm (Actiware v3.1) set to a medium sleep-wake threshold (< 40 counts⋅min^-1^ scored sleep) was then used to classify sleep and wake states. This threshold has demonstrated 87.8% agreement (i.e., percentage of sleep and wake epochs correctly detected) with polysomnography (29). The sleep diary was administered electronically and required participants to record their bedtime and get-up time. These data were used to verify bed- and get-up times determined via actigraphy (30). Measures obtained from actigraphy included sleep duration (total time spent sleeping) and sleep efficiency (sleep duration as a percentage of time-in-bed between bedtime and get-up time). Naps were not permitted throughout either condition period.

### Resistance exercise

The resistance exercise sessions were designed to stimulate muscle protein turnover and are described in detail elsewhere (21). Briefly, participants performed a standardised warm up followed by a 45-min resistance exercise session in groups of two or more participants. Each session encompassed six multi-joint exercises (barbell back squat, deadlift, leg press, bench press, seated cable row, lat-pulldown). Sets and repetitions were prescribed, and the load lifted (a percentage of 1-repetition maximum) increased each set from 60% (set 1) to 85% (set 4) of participant’s 1-repetition maximum.

### Hormone sampling and analysis

Venous blood samples were taken to assess estrogen and progesterone concentrations. Samples were immediately centrifuged for 15 min at 13,000 rev·min^-1^ at 4°C and the supernatant stored at -80°C prior to analysis. Estrogen and progesterone were each measured using automated competitive binding immunoenzymatic assays (Beckman Coulter, Sydney, NSW) (21).

### RNA extraction and quantification

RNA was extracted from pre-exercise muscle biopsies collected on day three and nine using an Allprep DNA/RNA/miRNA Universal extraction kit (Qiagen, Clayton, VIC) according to manufacturer’s instructions. Frozen muscle tissue (10-15 mg) was combined with 600 µL Buffer RLT plus (Qiagen) and homogenised with 650-800 mg silica beads in a MagNA lyser for two 30 sec homogenisation steps at 6500 rpm (Roche Diagnostics, North Ryde, NSW). The flow-through was then moved to the RNA column for RNA extraction, including a proteinase K and a DNase treatment steps according to the manufacturer’s protocol. A TapeStation System was used according to manufacturer’s instructions (Agilent Technologies, Mulgrave, VIC) to assess RNA quality and quantity, with an RNA integrity score of >7 considered acceptable.

### RNA sequencing and analysis

The RNAseq libraries were prepared using the Illumina TruSeq Stranded Total RNA with Ribo-Zero Gold protocol and sequenced with 150-bp paired-end reads on the Illumina Novaseq6000 (Macrogen Oceania Platform). Reads underwent a quality check with FastQC (v0.11.9), whilst Kallisto (v0.46.1) was used to map the reads to the human reference genome (HomoSapien GRCh38) and to generate transcript counts. Next, the dataset was normalised and filtered for low reads i.e., only genes with at least 10 reads on average were used in subsequent analysis. Genes with an average across all samples of 10 reads per million (RPM) or less reads (62%) were removed leaving a total of 15,543 transcripts for analysis. The analysis workflow is presented in supplementary material (Supplementary Figure 1). Differential expression analysis was performed using all RNA sequencing samples to compare NS and SR conditions by day, as well as the same condition by day. The DESeq R package (v1.40.2) (31) and pipeline were used to determine differentially expressed genes (DEGs) with significance thresholds set with an adjusted p-value of < 0.05 and Log_2_ fold change > 1 and <-1. Volcano plots and heatmaps, demonstrated in Figure 2 and Figure 4, were used to visualise expression patterns of DEGs across different contrasts.

**Figure 2:**
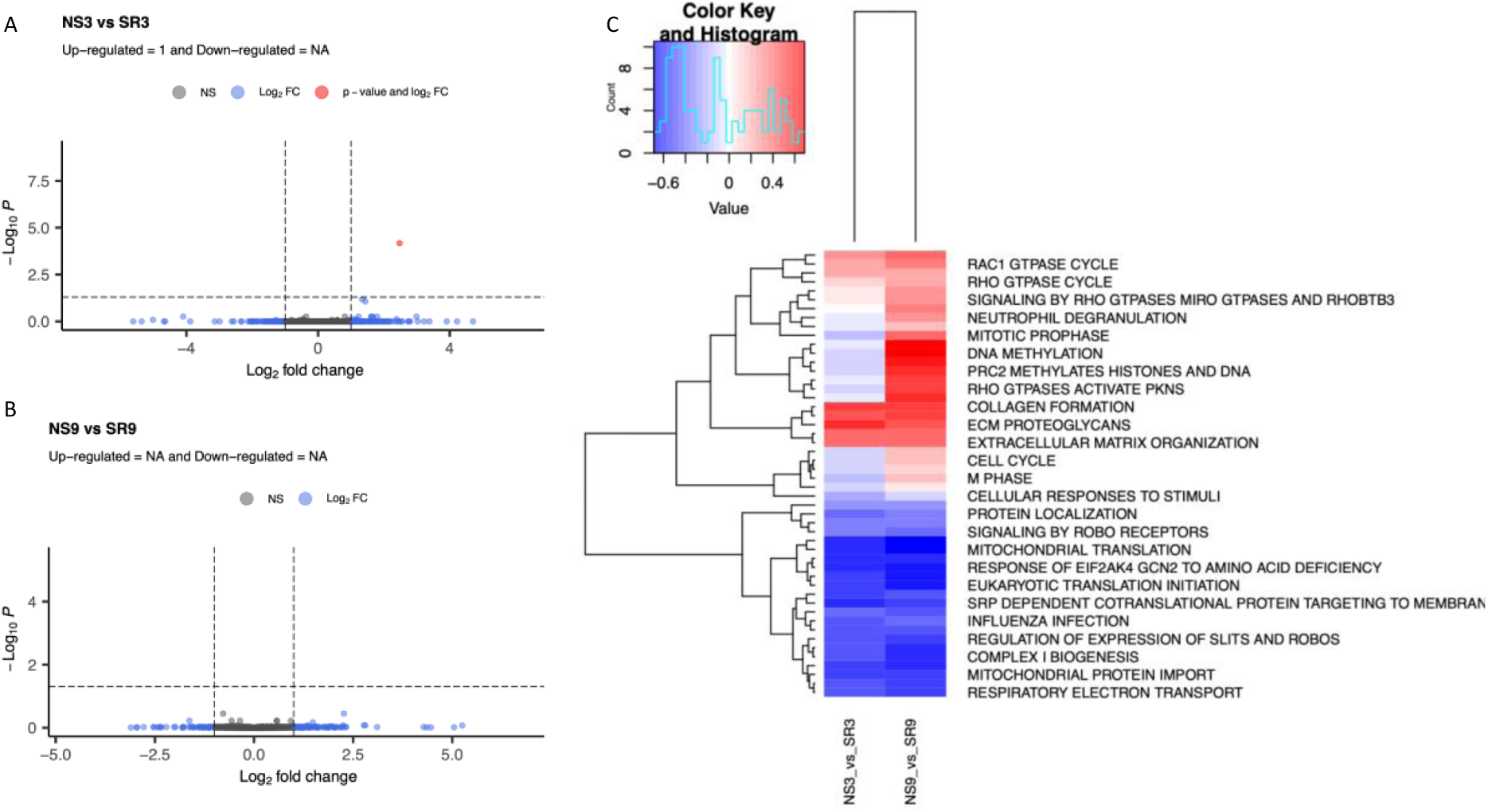
Volcano plots representing the differential expression of individual transcripts in the Normal Sleep (NS) vs Sleep Restriction (SR) condition in pre-exercise biopsies collected after three days **(A)** and nine days **(B)**. Blue dots represent values within the Log_2_ fold-change threshold (absolute value of > 1) between conditions. Red dots represent values within Log_2_ fold-change threshold and p-adjusted value threshold (p < 0.05) between conditions. Grey values are not within the Log_2_ fold-change and the p-adjusted value thresholds. Multi-contrast analysis **(C)** depicting the degree of up-regulation (red) or down-regulation (blue) of Reactome pathways in response to three (NS3_vs_SR3) or nine (NS9_vs_SR9) nights of sleep restriction when compared to three or nine nights of normal sleep, respectively.

**Figure 3.**
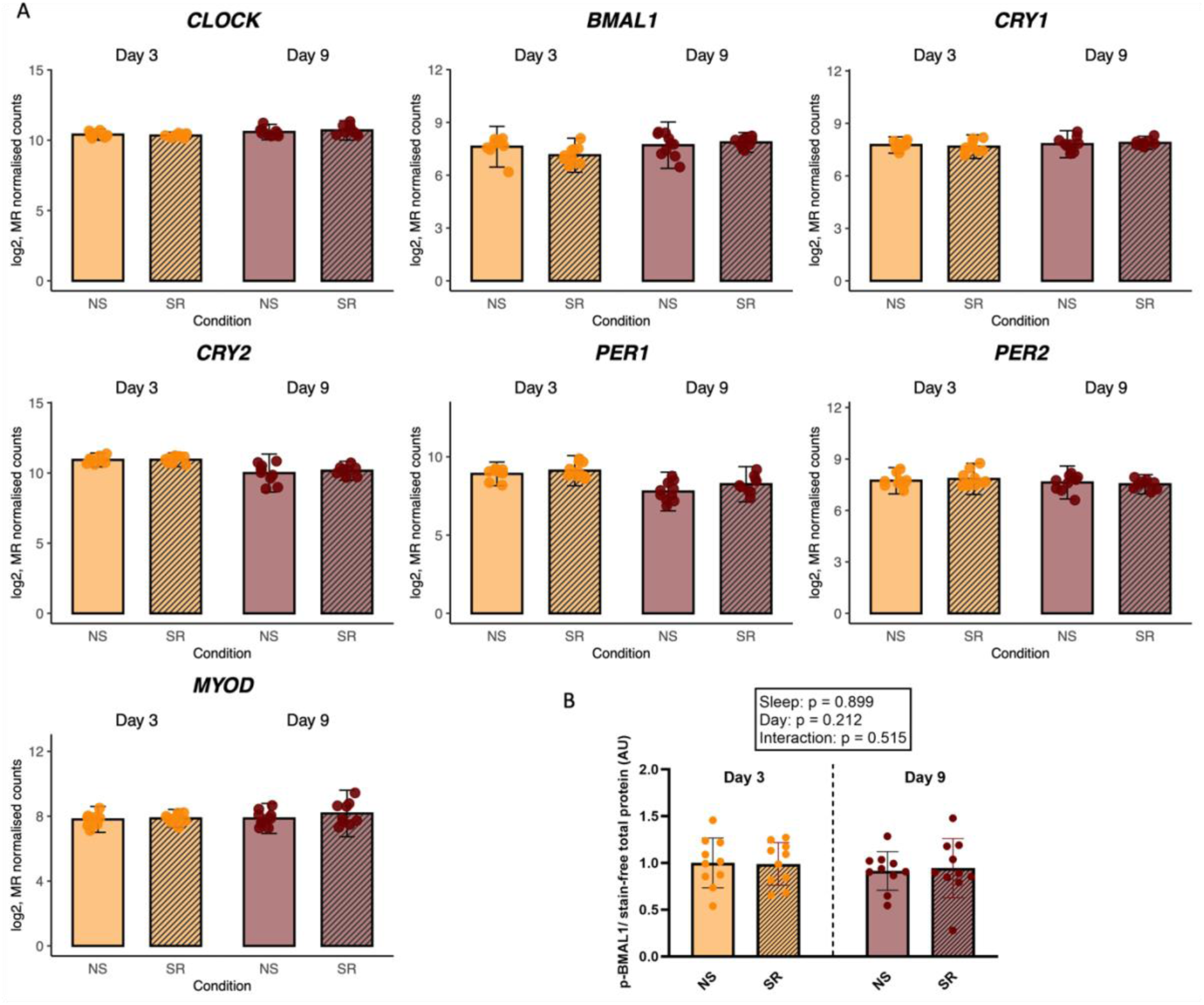
Gene expression of *CLOCK, BMAL1, CRY1, CRY2, PER1, PER2* and *MYOD1* **(A)** and protein expression of p-BMAL1 **(B)** in response to three or nine nights of sleep restriction when compared to normal sleep. Gene expression data were normalised using the median of ratios (MR) normalisation function from DESeq2. Protein expression data were normalised against total protein load and analysed using 2-ways ANOVAs. All Western Blots images are presented in Supplementary Figure 3a. NS = normal sleep; SR = sleep restriction. *P* < 0.05.

**Figure 4:**
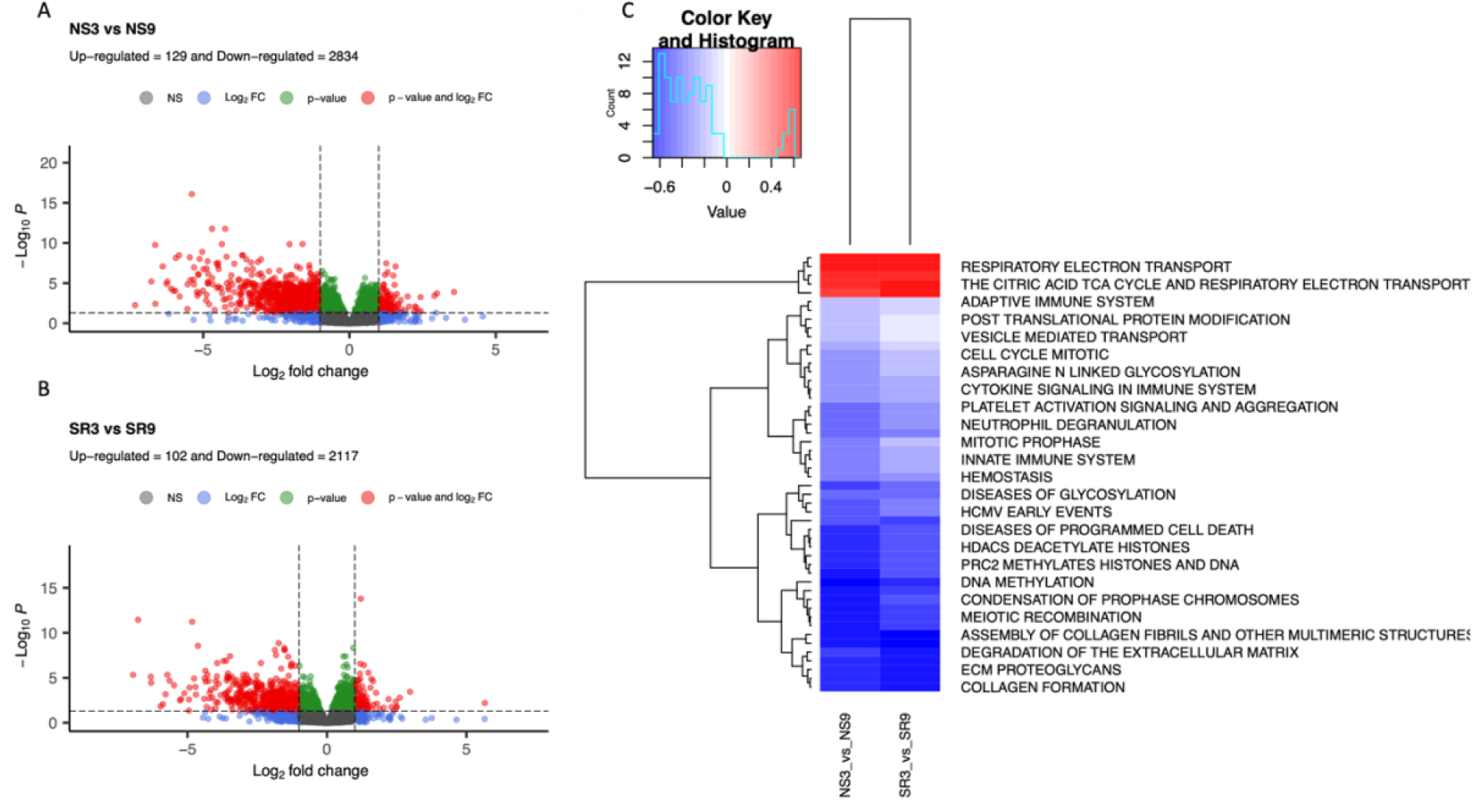
Volcano plots representing the differential expression of individual transcripts in pre-exercise biopsies collected following three sessions of exercise training in the control **(A)** and the sleep restricted condition **(B)**. Blue dots represent values within the Log_2_ fold-change threshold (absolute value of > 1) between conditions. Red dots represent values within Log_2_ fold-change threshold and p-adjusted value threshold (p < 0.05) between conditions. Grey values are not within the Log_2_ fold-change and the p-adjusted value thresholds. Multi-contrast enrichment analysis **(C)** depicting the up-regulation (red) or down-regulation (blue) of Reactome pathways in response to three resistance exercise training sessions in normal sleep (NS3_vs_NS9) or sleep-restricted (SR3_vs_SR9) conditions, respectively.

### Gene enrichment analysis

Gene enrichment analysis (GSEA) was performed to compare the transcriptomic response across different contrasts (i.e., SR and NS at day three and day nine). The analysis was performed using the Reactome pathways taken from the msigdbr package for the Homo Sapiens species. The GSEA was performed using the fgsea R package (v1.26.0) (32) using the four contrasts from the DESeq results as an input. The log_2_ fold changes were used for ranking genes and significance threshold was set to an adjusted p-value of <0.05. The Normalised Enrichment Score (NES) was used to plot the top 30 enriched pathways, which can be found in Figure 2, Figure 4 and Figure 5.

**Figure 5:**
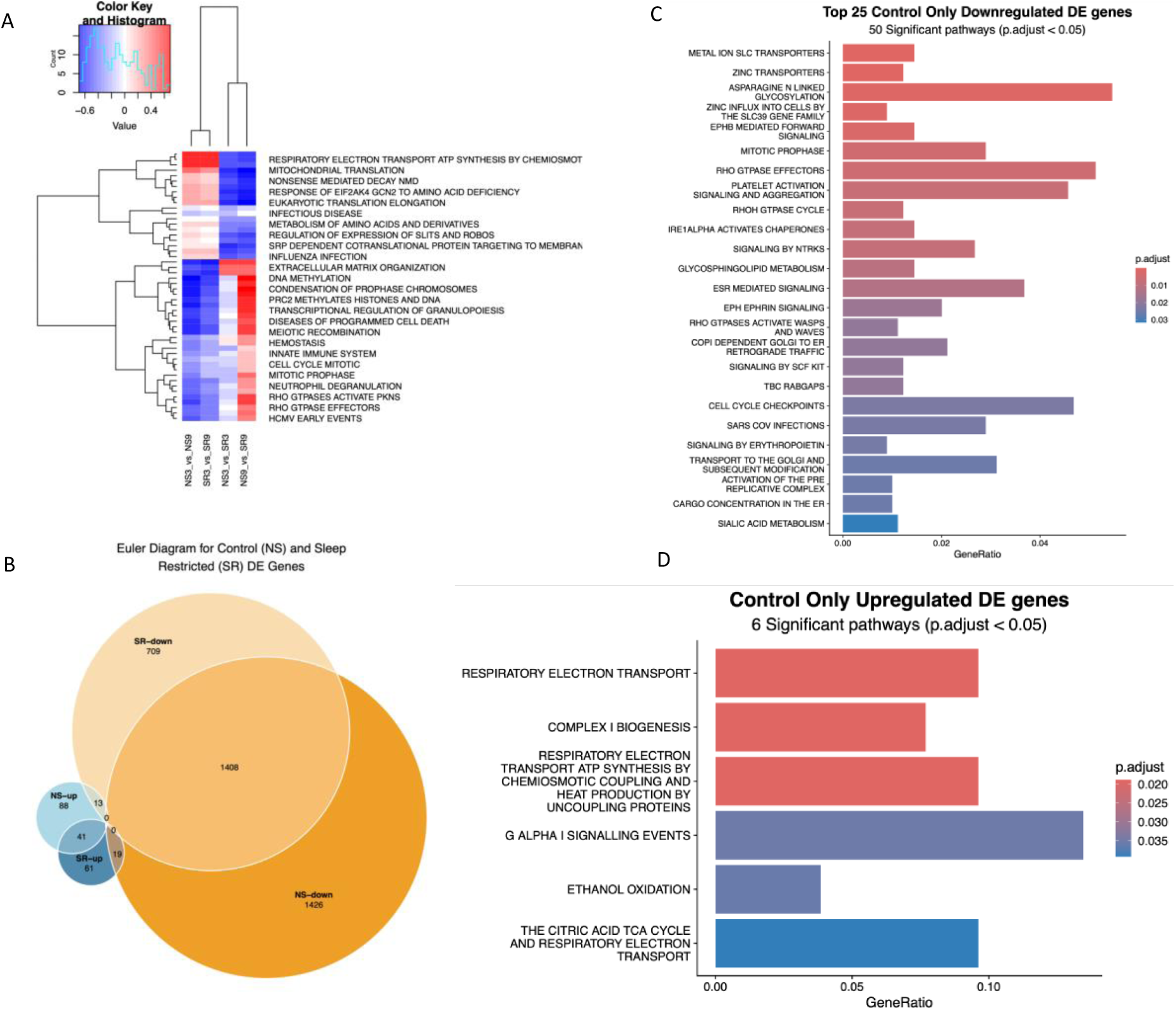
**A)** Multi-contrast enrichment analysis depicting the up-regulation (red) or down-regulation (blue) of Reactome pathways for all four possible contrasts. **B)** Euler diagrams depicting the number of genes differentially up-regulated (blue) and down-regulated (yellow) in response to three sessions of exercise training in the NS and SR conditions. **C)** Top-25 (of 50) Reactome pathways exclusively downregulated in the NS condition. **D)** Reactome pathways exclusively upregulated in the NS condition. FDR < 0.05.

### Multi-contrast enrichment analysis

Multi-contrast analysis was performed by contrast group (i.e., for NS vs SR and for same condition by day) using Mitch R Package (v1.12.0) (33). A second multi-contrast analysis was performed for all contrasts using the same package.

### Protein extraction and western blotting

Protein from pre- and post-exercise muscle biopsies collected on day 3 and 9 was extracted from approximately 25 mg of muscle using manual homogenisation in a RIPA buffer including phosphatase and protease inhibitors (Millipore, North Ryde, NSW). Total protein content was assessed using a BCA Protein Assay Kit (Thermo Fisher Scientific) according to manufacturer instructions. Following protein extraction, ∼ 20 µg of total protein from each sample mixed with reducing loading buffer (4x Laemmli buffer with 10% v/v 2-mercaptoethanol) were loaded into a 4–15% gradient Criterion Tris-Glycine extended (TGX) Stain Free gel (BioRad, Gladesville, NSW) and separated via electrophoresis at 120V for 60 min or 200 V for 40 min. The stain-free gel was imaged for total protein on a Universal Hood II GelDoc (BioRad, Gladesville, NSW) using the ImageLab v6 software (BioRad, Gladesville, NSW). Separated proteins were transferred to a methanol-saturated polyvinylidene difluoride (PVDF) membrane (Immobilon FL 0.45 µm #IPFL00010, Millipore, Billerica, MA) via wet transfer at 100 V for 45 min or using the Turbo Transfer system (BioRad, Gladesville, NSW).

For Akt (Cell Signalling Technology #9272), p-mTORC1 (Cell Signalling Technology #5536) and mTOR C1 (Cell Signalling Technology #2972), the membranes were blocked for one hour in 5% skim milk in Tris buffered saline plus 0.1% Tween-20 (TBST). Membranes were then incubated overnight at 4°C in primary antibodies diluted in Licor Intercept PBS blocking buffer (Licor, Lincoln, NE, USA) at a concentration of 1:1000 and washed in TBST. For the remaining antibodies, the membranes were washed in PBS, dried for 1 hour, re-saturated in methanol, washed in PBS and blocked for 1 hour in Licor blocking buffer. Membranes were then incubated in primary antibody diluted 1:1000 in blocking buffer with 0.2% v/v Tween-20 overnight at 4°C as follows: phospho-Akt Ser473 (Cell Signalling Technology #4060), p-4E-BP1 Thr37/46 (Cell Signalling Technology #9459), CLOCK (Cell Signalling Technology #5157), BMAL1 (Cell Signalling Technology #14020), and phospho-BMAL Ser42 (Cell Signalling Technology #13936) and washed in PBST. Membranes were then incubated with anti-rabbit or anti-mouse IgG Daylight® 680 nm (5366S, 5470S) or 800 nm (5151S, 5257S) secondary antibodies (all from Cell Signalling Technology) diluted 1:10,000 in Licor blocking buffer containing 0.2% Tween-20 and 0.01% SDS for 1 hour at room temperature.

Images were acquired using the Odyssey® Infrared Imaging System (Licor, Lincoln, NE) and blot densitometry assessed using the Odyssey v2.1 software (Licor, Lincoln, NE). Blot density and stain-free total protein density for each sample was calculated by linear regression using the standard curve constructed from a pooled sample loaded on each gel. Calculated blot density values were then expressed relative to the corresponding stain-free total protein density (34). The expression levels of CLOCK, BMAL1 and MYOD1 could not be reliably measured using either of the methods outlined above.

### Statistical analysis

Mean and standard deviation (SD) were calculated for all variables using the Stata statistical software (v17.0). Linear mixed models, with random intercepts for participants, were fitted for each outcome variable (protein expression, gene expression). The mixed models included main effects of sleep condition and day (for circadian variables, assessed pre-exercise only) or sleep condition, day and exercise (for MPS variables), and their interaction, with statistical significance set at p < 0.05. To account for any potential effect of the menstrual cycle on circadian gene and protein expression outcome variables (35), the concentration of estrogen and progesterone were tested as covariates in these models. In line with the most recent evidence suggesting minimal variation in the expression of exercise-regulated genes and proteins across the menstrual cycle (36, 37) ovarian hormones as covariates did not improve the fit of any of the tested models and were therefore not used. All values are expressed relative to the mean of the control variable (i.e., NS, day 3, pre-exercise) unless stated otherwise.

## Results

### Participants

The participants were aged 24.3 ± 4.8 yr with a body mass index of 23.6 ± 2.8 kg·m^-2^. There was no difference in mean daily protein intake (P = 0.98) or protein intake as a percentage of total macronutrients (P = 0.97) between conditions. Details regarding the physical strength and menstrual cycle of participants can be found elsewhere (21).

### Sleep

A detailed description and visual representation of the sleep duration, sleep efficiency and sleep quality results can be found elsewhere (21). Briefly, sleep duration was significantly reduced (SR, 4.7 ± 0.2 h; CON, 7.3 ± 0.8 h; *p* < 0.0005), and sleep efficiency significantly increased (SR, 95.0 ± 3.2%; CON 89.5 ± 5.6%; *p* < 0.0005) with SR. There was no change in sleep quality with SR.

### Effect of sleep restriction on the skeletal muscle transcriptome and circadian related protein expression

Whole transcriptome sequencing was performed on pre-exercise muscle biopsies collected on day 3 and 9. Three or nine nights of SR had no significant effect on the muscle transcriptome, independently of exercise. Of the 15,543 transcripts detected in skeletal muscle, only one (gametogenetin binding protein 2) was upregulated (FDR < 0.05) following three nights of SR when compared to control with no exercise (Fig. 2A). Similarly, no transcript was differently expressed at significance level following nine nights of SR when compared to normal sleep, in participants having undertaken the same exercise training regime at day 3, 5 and 7 (Fig. 2B).

Despite the lack of significant change in individual gene expression, we next investigated whether broader cellular pathways may be regulated in response to three or nine nights of SR. The top-15 Reactome pathways that were significantly up- or down-regulated at each time point are presented in supplementary material (Supplementary Figure 2a-b). We then used multi-contrast analysis to highlight the pathways that were regulated in the same or in opposite directions following three or nine nights of SR (Fig. 2C).

Cellular pathways that were downregulated in response to three and nine nights of SR included pathways related to translation (including rRNA processing (R-HSA-72312), eukaryotic translation elongation (R-HSA-156842), eukaryotic translation initiation (R-HSA-72613), translation (R-HSA-72766) and mitochondrial translation (R-HSA-5368287)) and oxidative metabolism (including respiratory electron transport (R-HSA-611105), complex 1 biogenesis (R-HSA-6799198) and the citric acid cycle (R-HSA-71403)). Cellular pathways that were upregulated in response to three and nine nights of sleep restriction pertained to extracellular matrix and cytoskeleton organisation (including collagen formation (R-HSA-1474290), integrin cell surface interaction (R-HSA-216083), ECM proteoglycans (R-HSA-3000178), degradation of the extra-cellular matrix (R-HSA-1474228), and extra-cellular matrix organisation (R-HSA-1474244)) as well as GTPases cycle and signalling. Of interest, several Reactome pathways were differentially regulated with time, meaning that they were downregulated as an initial response to three nights of SR but upregulated after nine nights. These included several cellular signalling pathways and cell cycle related pathways.

To verify our next hypothesis that circadian markers would be influenced by SR at the protein level, we next attempted to investigate the protein expression of known markers of circadian regulation in the same samples. Of circadian markers CLOCK, BMAL1, p-BMAL1 and MYOD1 it was only possible to obtain a robust and specific Western Blot signal for p-BMAL1 (Fig. 3B), which, in line with our gene expression analysis of core clock genes (*CLOCK, BMAL1, PER1, PER2, CRY1, CRY2, and MYOD1,* Fig. 3A), did not differ in response to three or nine days of sleep restriction, independently of exercise.

*Long-lasting effects of three repeated resistance exercise sessions on the skeletal muscle transcriptome* In striking contrast with the hypothesis that informed our study design, we found that nearly 3000 transcripts were still differentially regulated (FDR < 0.05) between muscle biopsies collected after three nights of normal sleep (pre-exercise, day 3) and muscle biopsies collected after nine nights of normal sleep and three sessions of exercise training (pre-exercise, day 9) (up-regulated = 128; down-regulated = 2834; Fig 4A). Details for each transcript can be found in supplementary material (Supplementary Table 1). This was despite the last muscle biopsy (pre-exercise, day 9) being collected after 48 hours of complete rest and our participants being highly trained. These results indicate a flow-on effect of the three training sessions completed between day 3 and 7 on the muscle transcriptome that remained visible 48 hours after the last exercise session. Similarly, 102 transcripts were upregulated and 2117 transcripts were downregulated (FDR < 0.05) after three nights of SR when compared to nine nights of SR and exercise training (Fig 4B), indicating a similar effect of training in SR conditions. The associated Reactome pathways are displayed in supplementary material (Supplementary Figure 2c-d).

Multi-level enrichment analysis revealed a strong overlap between the exercise-driven pathways that were differentially regulated after three exercise sessions both in the normal sleep and SR conditions (Fig. 4C). Strongly upregulated in both conditions were pathways pertaining to oxidative metabolism, including complex 1 biogenesis, respiratory electron transport, ATP synthesis and the citric acid cycle. Downregulated in both conditions were several pathways governing the cell cycle and cell structure organisation.

### Effect of sleep on the skeletal muscle transcriptomic response to exercise

Next, multi-contrast analysis including all four possible contrasts indicated that sleep restriction and exercise had contrary effects on the skeletal muscle transcriptomic pathways (Fig. 5A). The Reactome pathways that were upregulated by exercise (typically oxidative metabolism and translation) were also the pathways that were downregulated by SR. In contrast, the pathways that were downregulated by exercise in both normal sleep and SR conditions (including some elements of cell cycle and cellular structure regulation), were upregulated after nine days of SR, but not after three days of SR.

Since we observed an unexpectedly strong effect of three training sessions on the muscle transcriptome despite the last biopsy having been collected after 48 h of rest, we next sought to understand whether this response to training was moderated (i.e. blunted or exacerbated) by SR. Euler diagrams were used to visualise whether the genes differentially regulated in response to three sessions of exercise training were the same in both the NS and SR conditions (Fig. 5B), where a larger overlap indicates a smaller mediating effect of SR. Of the 2117 and 2834 genes downregulated in the SR and control conditions, respectively, only 1408 were common to both. Similarly, of the 102 and 129 genes respectively upregulated in the SR and control conditions, only 41 were the same between both conditions, overall indicating an important moderating effect of sleep restriction on the response to exercise.

Over-representation analysis was used to investigate the nature of the exercise driven genes that were exclusively upregulated in the control (n=88) or SR (n=61) condition, or exclusively downregulated in the control (n=1426) or SR (n=709) condition. Fifty pathways including cell cycle components and a number of molecular signalling cascades were significantly represented by the exercise driven genes exclusively downregulated in the NS condition, while no pathway was significantly represented by the genes exclusively downregulated in the SR condition (Fig. 5C). Similarly, six pathways, including once again components of oxidative metabolism and mitochondrial respiration, were significantly represented by the genes exclusively upregulated in the NS condition, while no pathway was significantly represented by the genes exclusively upregulated in the SR condition (Fig. 5D). This confirms that SR mediates the skeletal muscle response by blunting the metabolic and cellular adaptation to resistance exercise training.

Finally, an acute increase in muscle protein synthesis in the 1-3 hours that follow the end of a session is a well-established response to resistance training (38). To examine the interaction of resistance exercise and SR on muscle protein synthesis activation, we assessed the protein levels of common markers of muscle protein synthesis using Western Blotting pre- and one hour post- the first (Day 3) and last (Day 9) exercise session. Linear mixed models only revealed a sleep × day × exercise interaction for p-mTORC1 (*p*=0.03), which did not translate into any significant, relevant pair-wise comparison (Fig. 6).

**Figure 6.**
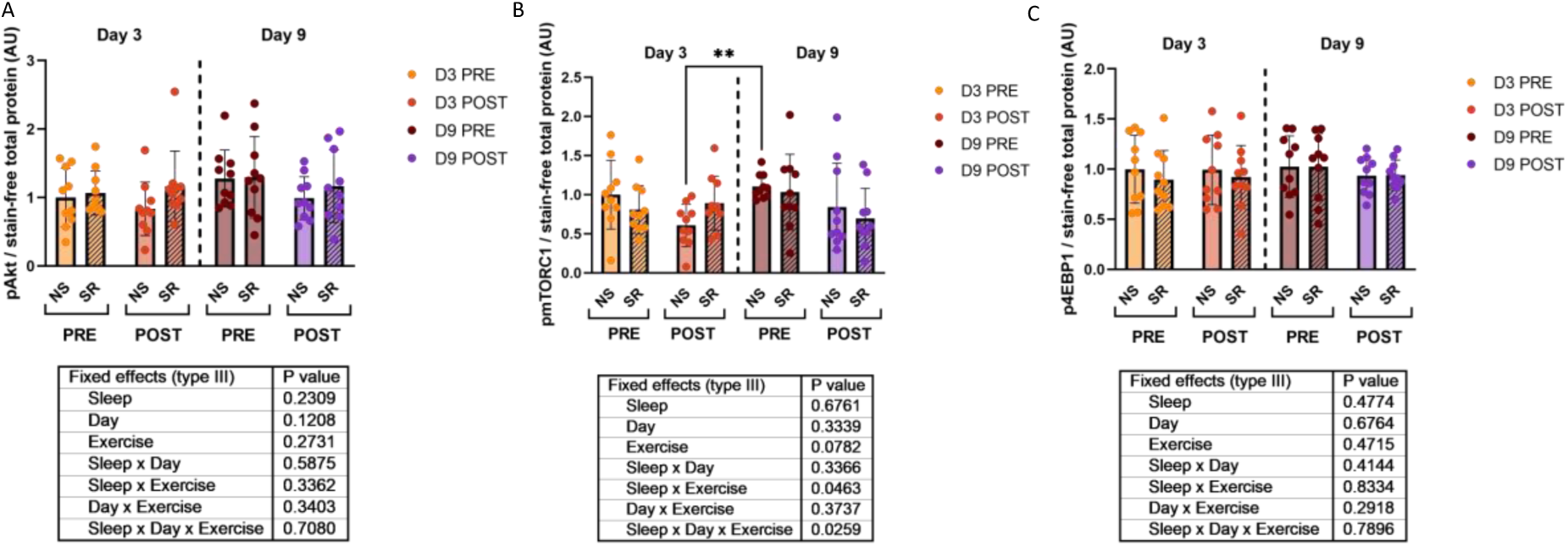
Pre- and post-exercise protein expression of p-Akt **(A)**, p-mTORC1 **(B)** and p-4EBP1 **(C)** in response to three or nine nights of sleep restriction when compared to normal sleep. Protein expression data were normalised against total protein load and analysed using linear mixed models. All Western Blots images are presented in Supplementary Figure 3b-d. *P* < 0.05.

## Discussion

Our study investigated the effect of nine nights of sleep restriction, with repeated resistance exercise sessions, on the skeletal muscle transcriptomic response in young, resistance trained females. Contrary to our hypothesis, we observed no significant effect of the sleep restriction protocol on differential gene expression after either three days of sleep restriction when compared to normal sleep, or nine days of sleep restriction combined with resistance exercise when compared to normal sleep combined with resistance exercise. Unexpectedly, despite muscle samples being collected 48 h after the last resistance exercise session, approximately 3000 genes were still differentially regulated at this time point in both the normal sleep and sleep restricted conditions. Pathway analysis followed by multi-contrast analyses suggested a blunting effect of sleep restriction on the transcriptional response to resistance exercise. These findings have practical implications for optimising the beneficial effects of exercise in populations that are commonly unable to obtain adequate sleep.

Skeletal muscle is a key metabolic organ that has its own internal biological clock. By negatively impacting the regulation of dozens of gene pathways, whole body and skeletal muscle specific metabolism (39, 40), sleep loss has significant detrimental effects on the organism. Here we report that both three and nine nights of sleep restriction did not cause significant changes to individual gene expression in skeletal muscle, with only one differentially expressed gene reported after three days of sleep restriction when compared to normal sleep. This gene, gametogenetin binding protein 2, is an angiogene that has not been characterized in skeletal muscle. This observation is contrary to the hypothesis we formulated at the time this study was designed, where a previous study had investigated the effect of one night of total sleep deprivation (24 h wakefulness) and reported 118 differentially expressed genes (DEGs) in skeletal muscle (41). However, our findings are now supported by a recent study that investigated the skeletal muscle transcriptomic response to a five-night period of sleep restriction (4 h time in bed per night) and also reported no differential gene expression (11). These discrepancies may stem from differences in the length or severity of the sleep loss interventions implemented. There is a well-documented ‘dose response’ effect to sleep loss in many aspects of physiology including cognitive function (42), glucose tolerance (39, 41), and vascular endothelial function (43), whereby the degree of detrimental effects depends on the extent of sleep loss, albeit in a non-linear manner. Despite the lack of differentially regulated individual transcripts, there are many similarities between the enriched transcriptional pathways reported in our study and the previous studies that have examined the skeletal muscle transcriptome responses to either one or five nights of sleep loss. Pathways associated with protein translation and oxidative metabolism were similarly down-regulated after one night of sleep deprivation (41) or five nights of sleep restriction (11) and support translational findings that skeletal muscle protein synthesis (6, 7) and mitochondrial respiratory function (44) are negatively influenced by sleep restriction. While there are large differences in the number of DEGs between acute sleep deprivation (i.e., 24 h wakefulness) and chronic sleep restriction (i.e., less than 7 h time in bed) studies (11, 41), there are similar enrichment profiles of transcriptional pathways, suggesting that the magnitude of change may be influenced by the length and severity of the sleep intervention.

While the enrichment of cellular pathways related to protein translation and oxidative metabolism were downregulated with three and nine nights of SR, a number of cellular signalling and cell cycle related pathways were differentially regulated with time (i.e., up-regulated after three nights of SR when compared to normal sleep, and then down-regulated after nine nights of SR when compared to normal sleep in the same exercise conditions). Since the effect of exercise is already accounted for in our contrasts (i.e. comparisons are only made between groups having performed the same amount of exercise), it is plausible that a compensatory or adaptive response to the stress associated with a sleep restriction intervention could have occurred. Allostasis refers to the physiological process that maintains homeostasis, recognising that set points and other boundaries of control may change with environmental conditions (45). Previous sleep research has demonstrated similar patterns of change whereby the greatest effect of sleep loss is observed after one night but improves or remains steady with subsequent nights of sleep restriction (46). For example, one study demonstrated that insulin sensitivity was reduced following one week of sleep restriction (1.5 h < habitual sleep duration), however, when the intervention was continued for two- and three-weeks, insulin sensitivity had returned to baseline (47), suggesting an adaptive response. How this concept fits within the strong epidemiological data that demonstrates significantly increased risk of all-cause mortality (48), cardiovascular disease (49) and type 2 diabetes (50) with sleep loss, however, requires further research, including in long-term sleep restriction conditions.

Resistance exercise is a potent stimulus for promoting positive adaptations in skeletal muscle size, strength, and function. Many of the adaptations induced by resistance exercise within skeletal muscle are underpinned by a distinct and dynamic transcriptomic response. There is an underlying assumption in the exercise physiology field, mostly informed by early studies, that the vast majority of differentially expressed genes have returned to baseline 48 hours post exercise. Our results however indicate that irrespective of sleep, ∼2000-3000 genes were still differentially expressed 48 h post the completion of three resistance exercise sessions. In muscle biopsies taken up to 24 h post resistance exercise, similarly large transcriptomic responses have previously been reported (20, 51, 52). The transient nature of the transcriptional response to resistance exercise, however, is not well characterised. Early studies by the Trappe group established that peak gene induction of myogenic genes occurred ∼4–24 h post exercise, while peak proteolytic gene induction occurred as early as 1– 4 h post exercise (53, 54). Some previous studies have also used aerobic exercise to demonstrate the variable time-course of gene expression in skeletal muscle. For example, Kuang et al. (55) highlighted that the temporal pattern of 22 genes following a single session of high-intensity interval exercise was highly variable, with the largest changes in some exercise-responsive genes occurring between 3 and 72 h post exercise. While the number of differentially expressed genes reported in the current study is consistent with previous transcriptomic responses 3-24 hours after a single resistance exercise session (47–49), our findings provide a novel addition to the temporal response to resistance exercise and suggest that a significant number of genes are still differentially expressed 48 h post-exercise.

Our findings further established that sleep and exercise have an opposing effect on the muscle transcriptome. Lin et al. (11) showed changes in the muscle transcriptomic response to repeated high intensity interval exercise with five nights of sleep restriction in males, but did not have a normal sleep and exercise group by which to specify the influence of sleep restriction on this response. Our study identified several thousands of genes that were differentially expressed with exercise, where only 39% of downregulated and 18% of upregulated genes were common between normal and restricted sleep. This suggests that the response to resistance exercise is blunted when individuals are exposed to sustained sleep restriction. Exercise performed during periods of sleep restriction mitigates a number of negative transcriptional changes, such as those involved with mitochondrial dysfunction (11) yet our findings suggest that performing exercise when sleep restricted may not provide the same adaptive response for individuals as if they were fully rested. Taken together with our group’s performance-based data whereby the quantity (i.e., volume load output) and quality (i.e., bar velocity) of lifting performance was reduced under the same conditions of sleep restriction (21), our data suggests a clear detriment to the skeletal muscle molecular and functional response when trained females are exposed to sustained sleep restriction. These findings may have broader implications for people performing resistance exercise during periods of inadequate sleep. For example, athletes or military personnel who undergo periods of sleep loss during training cycles (56, 57) may therefore have a diminished adaptive response that are critical for improvements in performance, reducing injury risk, and improving health outcomes.

When comparing our findings to the existing literature, two main study-specific limitations may be kept in mind. Firstly, differences may in part reflect the time of day and amount of time since waking that the muscle sample was collected. Indeed, biopsies in our study were collected 8 hours after waking, for the exercise session to be performed at the highest point of sleep propensity and prior to the second circadian peak in alertness during the afternoon (58, 59), whereas previous studies collected muscle samples shortly upon waking (11, 41). The timing of muscle sample collection may therefore have influenced the transcriptional response observed, which represents a snapshot of the skeletal muscle transcriptome at a specific time point and may potentially not reflect the transient nature of the changes in gene expression that occur with sleep restriction. In addition, there may be a sex-specific effect to consider given the majority of existing sleep and exercise studies have been performed in male participants (14, 15). Indeed, there is evidence that sex influences the skeletal muscle transcriptome in response to resistance exercise (60). A recent meta-analysis found 247 genes (pooled from 43 studies) that were differentially expressed according to sex in response to various exercise modes (61), which may account for some of the differences between this study and the existing literature.

In conclusion, both three and nine nights of sleep restriction (5 h per night) altered the enrichment of skeletal muscle transcriptomic pathways, albeit without detecting any differential expression of individual genes in young, resistance trained females. In contrast, performing three successive resistance exercise sessions under both normal sleep and sleep restricted conditions induced the differential expression of several thousand genes that remained altered 48 h after the previous exercise session. The transcriptional response to these successive bouts of resistance exercise was moderated by the sleep restriction condition, indicating that the physiological health and performance benefits associated with skeletal muscle transcriptional responses to resistance exercise may be reduced when performed under conditions of limited sleep. This may have implications for physically active populations performing resistance-based exercise under sleep-restricted conditions and their subsequent skeletal muscle training adaptation, and should encourage further research into the impact of exercise prescription on the muscle molecular response. Our findings should also prompt future studies to explore the role which sex, training status, and exercise and biopsy timing have on the muscle transcriptome with resistance exercise.

